# Mode of Growth and Temperature Dependence on Expression of Atrazine-degrading Genes in *Pseudomonas* sp. strain ADP Biofilms

**DOI:** 10.1101/302877

**Authors:** Michael A. Delcau, Victoria A. Henry, Emily R. Pattee, Tonya L. Peeples

## Abstract

Bacterial strain *Pseudomonas* sp. strain ADP is capable of metabolizing atrazine, a synthetic herbicide, and uses atrazine as a sole nitrogen source for growth. The microbe completely mineralizes the substrate in a catabolic pathway comprised of six enzymatic steps. All enzymes, AtzA-AtzF, encoded by corresponding genes, *AtzA-AtzF*, are located on a self-transmissible plasmid, pADP-1. (Souza, M. L., Wackett, L.P., and Sadowsky, M.J *Appl. and Environ. Microbiol*. 64(6): 2323-2326, 1998) RT-qPCR was used to differentiate gene expression in atrazine-degrading genes in *Pseudomonas* sp. strain ADP cells grown as suspended cells and as biofilms. Relative gene expression was also evaluated for biofilms grown at 25°C, 30°C, and 37°C. Complementary atrazine kinetic data was collected using GC-MS for both modes of growth and temperature variance. No significant difference in expression was observed for all atrazine-degrading genes in biofilm-mediated cells relative to planktonic cells, suggesting neither decreased or increased catabolic activity at the mRNA level. In contrasting experiments concerning biofilm growth, expression was downregulated at 37°C for genes AtzA, *AtzB*, and *AtzC* and upregulated for genes *AtzD, AtzE, AtzF*, signifying *Pseudomonas* sp. strain ADP biofilms catabolic activity may change in response to substantial temperature changes. Gradual decreases in atrazine concentration were apparent in cells grown in shake flasks, while biofilm-mediated cells showed transient increases and decreases in reactor effluent. The complex extracellular matrix components, quorum sensing, and genetic transfer may account for accumulation and rapid degradation of atrazine. The data collected suggest biofilm-mediated bioremediation may give insight into catabolic activity and atrazine degradation potential.

**Importance:** Atrazine is the second most applied herbicide in the United States. It is applied to crops including sorghum, corn, and sugarcane to prevent the growth of broad-leaved weeds. Once used, it can permeate the soil and contaminate proximal groundwater sources, which provide drinking water for over 90-million people. The Environmental Protection Agency sets the maximum contaminant level at 3 parts per billion for atrazine in drinking water, however this is frequently exceeded in rural regions which presents a public safety concern. Atrazine is an endocrine disruptor compound and a suspected teratogen in humans and freshwater species, respectively. This research is significant in evaluating the use an atrazine-degrading strain, *Pseudomonas* sp. strain ADP, grown in a biofilm mode of growth to increase the degradation potential compared to suspended cells. Our results concerning expression and kinetics will aid the development of biofilm reactors for *ex situ* bioremediation and understanding environmental biofilms.

## Introduction

Atrazine is a ubiquitous herbicide used to control the pre- and post-emergence of broadleaved weeds in crops such as maize, sugarcane, and sorghum. It is one of several chemicals, including simazine and propazine, that falls under the triazine class of herbicides due to their similar mechanisms of toxicity in weeds.(1) Structurally, atrazine is a nitrogenous ring with chloride, ethylamine, and isopropylamine moieties. The synthetic molecule is an odorless, white powder in its pure form.(2) Despite contentious evidence surrounding the molecule’s carcinogenicity, atrazine has demonstrated endocrine-disrupting effects on aquatic fauna at higher concentrations from runoff into natural water systems.(3–5)

The chemical has been banned from use in the European Union due to unpreventable water contamination since 2003, however in the U.S. it is one of the most used pesticides in agriculture, only second to glyphosate.(6, 7) The Environmental Protection Agency has restricted the acceptable amount of atrazine in drinking water by instituting a maximum contaminant level (MCL) of three parts per billion. Despite the enforced limit, some areas notably in the Midwest, have demonstrated detected atrazine concentrations exceeding this level in raw drinking water samples.(8)

Facilitating degradation of these compounds is therefore vital for pollutant-free soil and groundwater. One approach, termed bioremediation, depends on optimal environmental conditions and microbial metabolism to break down persistent pollutants such as petroleum hydrocarbons in the ocean, heavy metal contaminants in drinking water, and polyaromatic hydrocarbons. Two approaches, biostimulation and bioaugmentation, rely on support of microbial growth though limiting nutrients and the addition of cells to a site for degradation. In order to optimize remediation - contaminant bioavailability, moisture content, oxygen content, and temperature at the site of pollution are to be considered.(9) *Pseudomonas* sp. strain ADP, was isolated and identified as an organism capable of mineralizing s-triazine herbicides, including atrazine.(10)

Mandelbaum demonstrated *Pseudomonas* sp. strain ADP is capable of metabolizing atrazine into carbon dioxide and ammonia.(10) The first three enzymatic steps encoding for the genes *AtzA, AtzB*, and *AtzC*, follow an initial hydrolytic dechlorination of atrazine to yield hydroxyatrazine, and subsequently two deamidations to form n-isoprolylammelide and cyanuric acid, respectively.(11) Enzymes AtzD, AtzE, and AtzF, complete the catabolic steps by several biotransformations until the substrate is completely mineralized to carbon dioxide and ammonia(12, 13) (FIG 1). All six enzymes in the pathway are encoded by genes on a single 108kb plasmid, pADP-1. Genes AtzA, *AtzB*, and *AtzC* are dispersed throughout the plasmid while *AtzD, AtzE*, and *AtzF* are contiguously clustered in an operon-like manner.(12) The catabolic capacity of strain *Pseudomonas* sp. strain ADP has been extensively studied while characterizing cell growth on agar plates, suspended in media, and dispersed in soil.(14–16) However, to our knowledge, this strain has solely been considered in a sessile lifestyle as a biofilm in kinetic, Raman, and spherical stirred tank reactor experiments.(17, 18)

**FIG 1.**
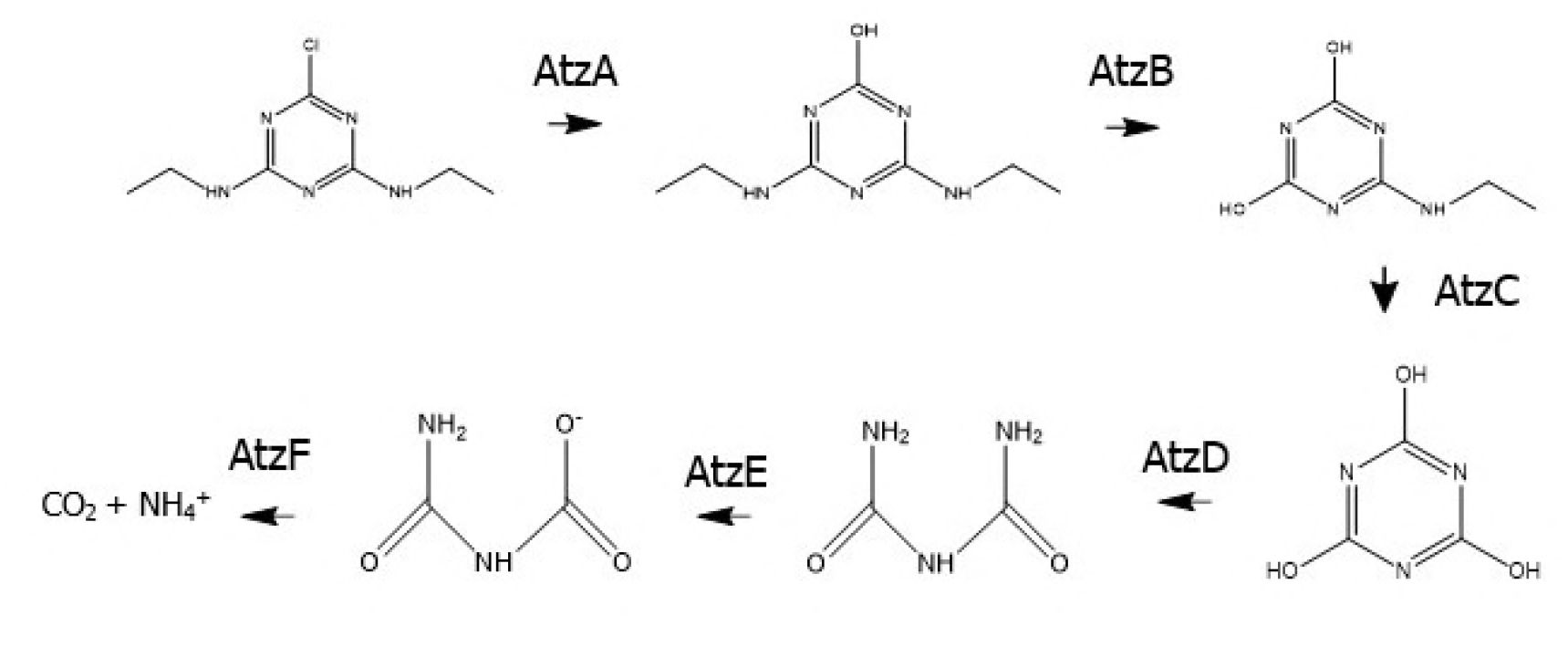
Stepwise catabolic reaction taken by *Pseudomonas* sp. strain ADP cells to degrade atrazine by the activity of six enzymes.

Biofilms are formed by the assembly of microbes from one single or multiple species that attach to an abiotic or biotic surface. These films are characterized by the secretion of extracellular polymeric substances to form an encompassing matrix.(19) This results in phenotype alternation and changes in gene expression and cellular growth. Besides being a more common mode of growth for bacteria in natural environments, employing biofilms in bioremediation yields several advantages. Some of these include protection from outside forces and/or stressors, increased genetic exchange and communication, and greater nutrient availability.(20, 21) Biofilms have been successfully used in bioremediation concerning persistent organic pollutants (POPS), polycyclic aromatic hydrocarbons (PAHs), and dioxins.(22–24)

The genetic activity of microbes during degradation of atrazine may enable further insight into the efficiency of microbial metabolism.(25) Devers demonstrated that catabolic activity of the atrazine-degrading genes in *Pseudomonas* sp. strain ADP began immediately once the herbicide was added. The entire *atz* gene set was either basally expressed or upregulated in response to low- and high levels of atrazine treatment.(13) While Devers reported expression patterns for atrazine catabolism in free cells, to date there is little published information regarding the expression of genes which encode atrazine degradation for cells cultivated as biofilms.

The kinetic parameters of *Pseudomonas* sp. strain ADP growth and atrazine degradation have been measured under variance of stirring speed and temperature in both shake flasks and bioreactors.(17) Most recently, Raman spectroscopy has been used as a novel tool to differentiate between *Pseudomonas* sp. strain ADP planktonic cells and biofilms, largely due to the presence of a characteristic extracellular polymeric matrix present in the resulting spectra.(18) Our goal in this work is to provide gene expression data as it relates specifically to biofilms in atrazine bioremediation, as this may aid in evaluating our hypothesis concerning *Pseudomonas* sp. strain ADP biofilms demonstrating increased catabolic activity at the mRNA level compared to suspended cells. The gene expression studies in this current work have been performed while evaluating a mode of growth and temperature dependence model in addition to corroborative analytical chemistry experiments.

## Results

A growth curve of *Pseudomonas* sp. strain ADP cells using mineral medium containing atrazine (30 mg L^-1^) was obtained to ensure harvest of high cell density for biofilm growth (FIG 3). It was determined cells could be collected for biofilm growth at 72 hours prior to centrifugation and re-suspension, optical density at 600 nm (OD_600_) of 0.3. The primers used to assay for each corresponding gene are listed in Table 1, with 16s rRNA used as a validated reference gene.(13)

**FIG 2.**
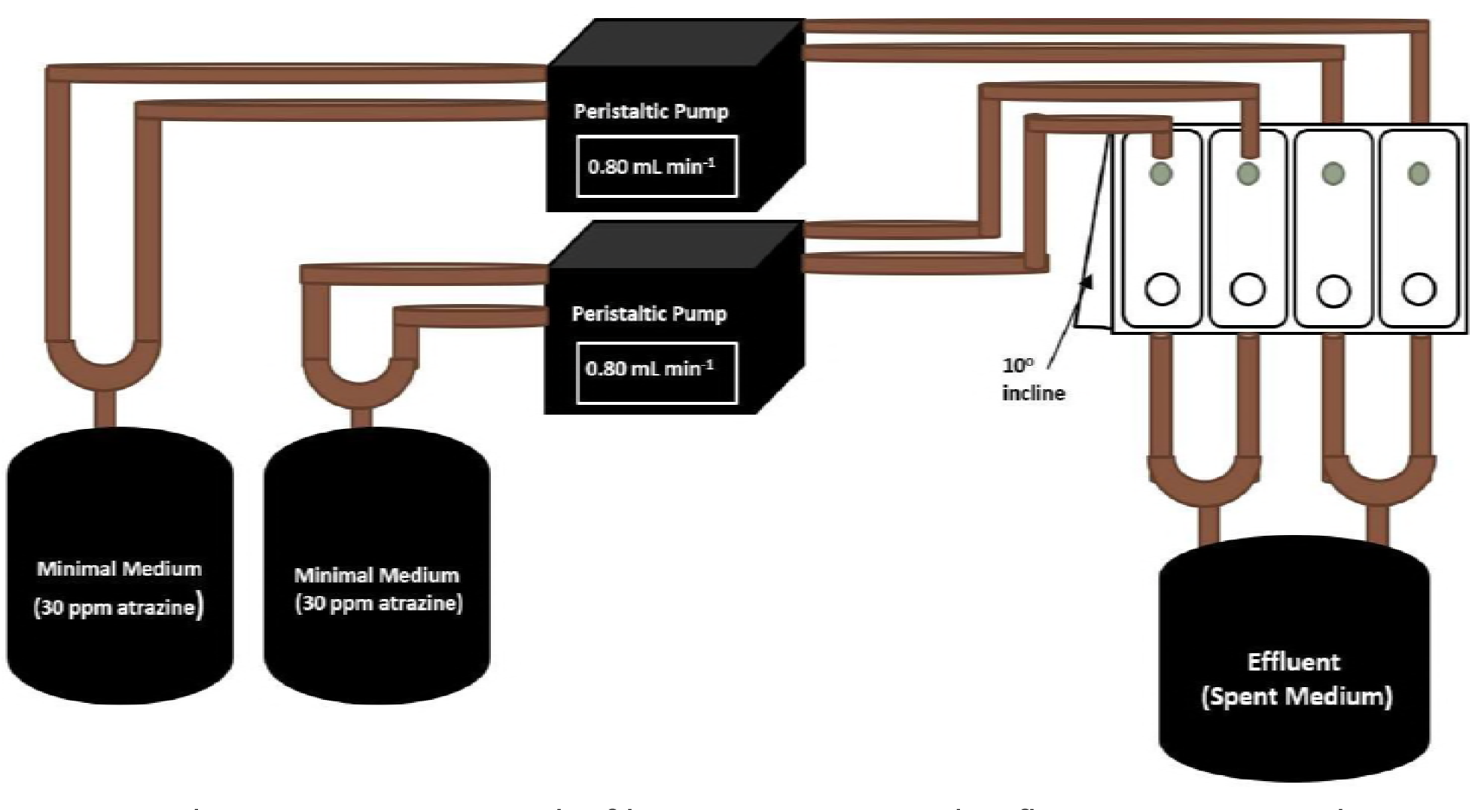
*Pseudomonas* sp. strain ADP biofilms were grown in a drip-flow reactor. Minimal medium containing atrazine was propelled via peristaltic pumps to a four-chamber reaction vessel containing Silane-coated slides at a 10-degree incline.

**FIG 3.**
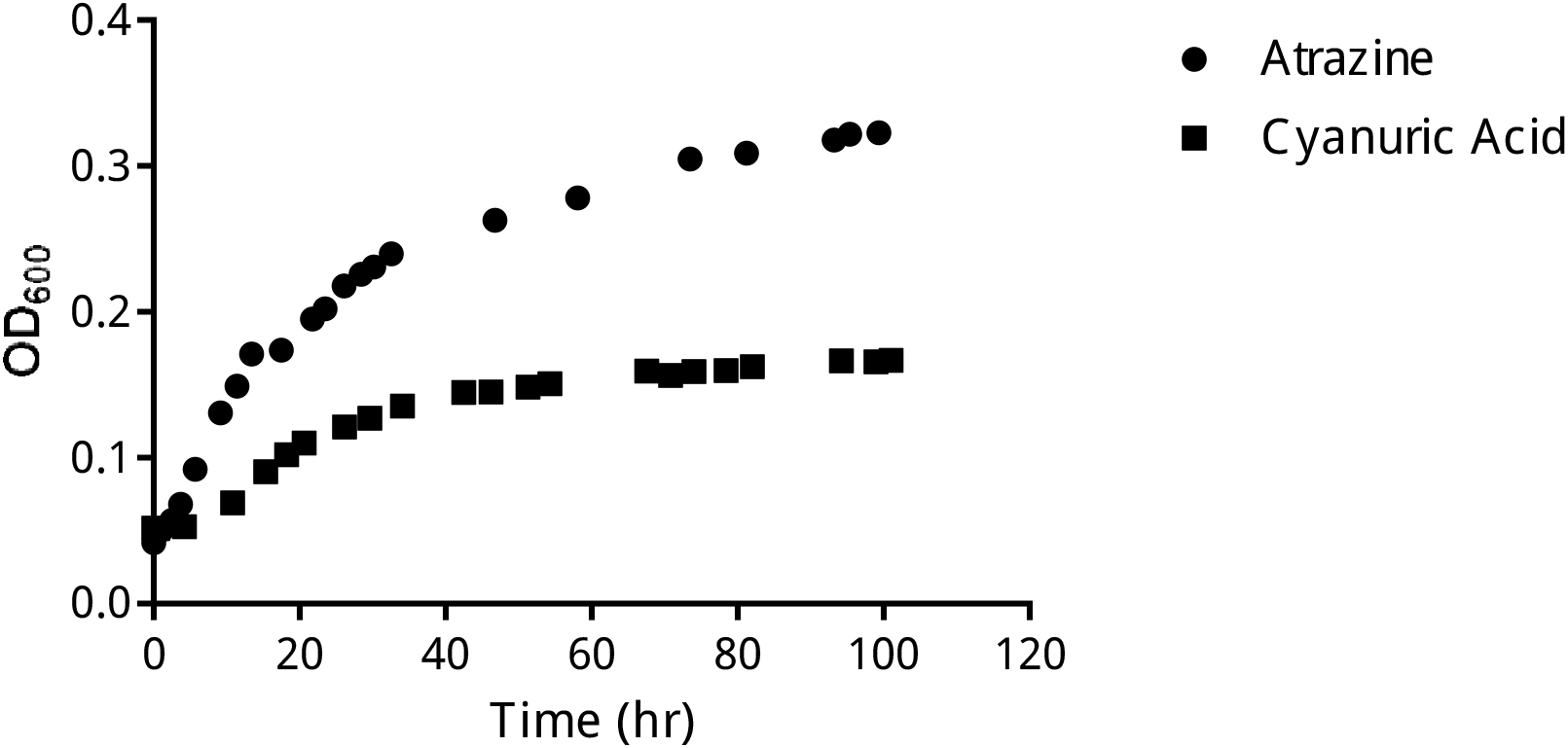
Growth of *Pseudomonas* sp. strain ADP under minimal medium containing 30 ppm of atrazine as a sole nitrogen source and cyanuric acid as a sole nitrogen source.

**TABLE 1.**
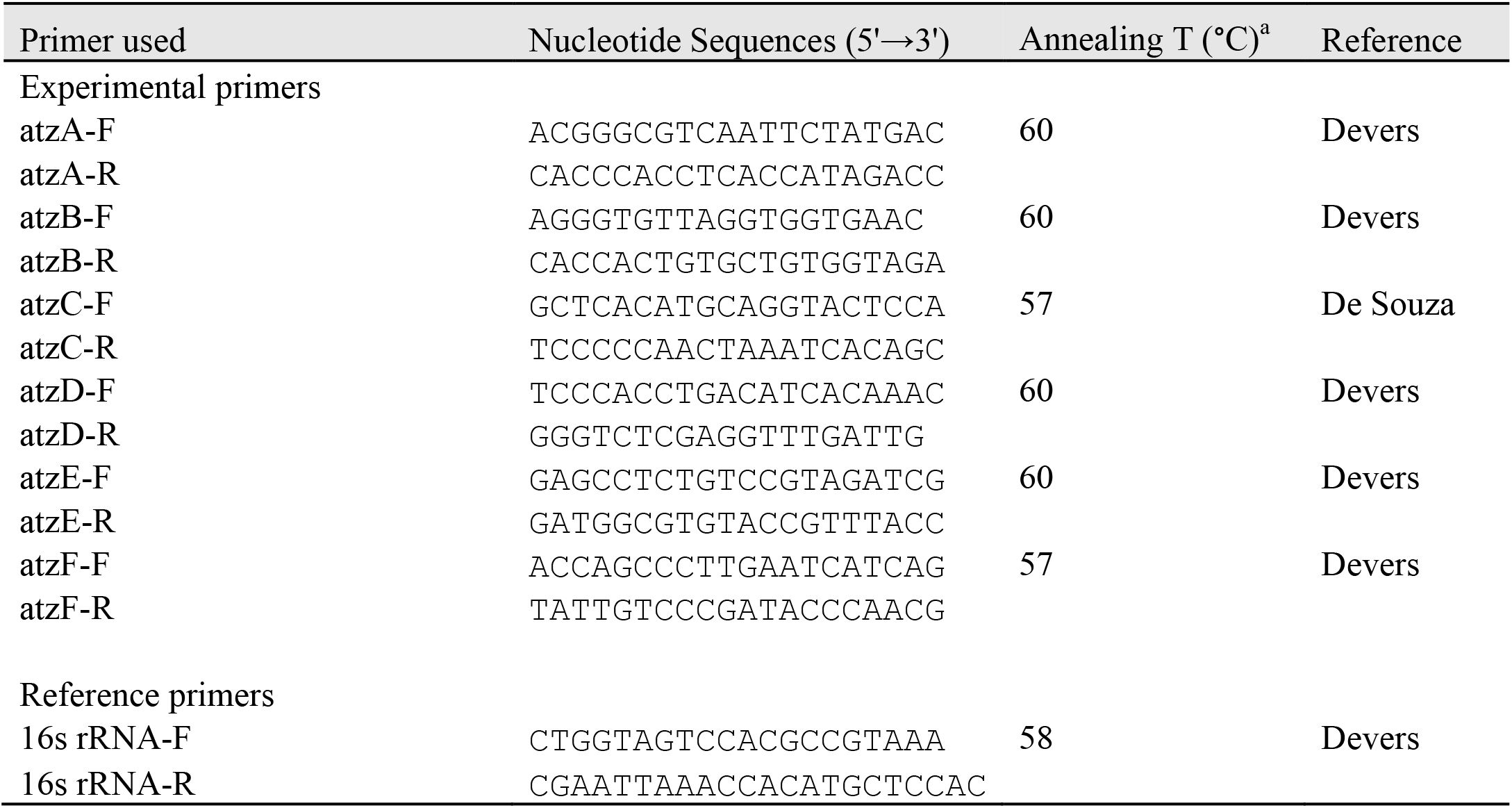
Primers used in this study for gene expression

644

Cyanuric acid, a metabolite of atrazine degradation, was used as an alternative sole nitrogen source for *Pseudomonas* sp. strain ADP growth and metabolism. Once growth was validated via streak plates of bacterial cells on plates containing mineral medium (500 mg L^-1^ cyanuric acid) with evident clearing zones and microbe proliferation, a similar growth curve was conducted (FIG 3) over a period of approximately 100 hours. A short lag phase, log phase, and beginning of stationary phase are demonstrated from the start to end of the growth period. The growth rate of *Pseudomonas* sp. strain ADP cells in liquid mineral medium containing cyanuric acid (OD_600_ of 0.10 at t = 20 hours) is approximately one-half the growth of rate of cells in medium containing atrazine (OD_600_ of 0.18 at t = 20 hours) in exponential phase.

The relative gene expression marginally decreased in biofilm cells compared to cells derived from shake flasks for gene *AtzA*. The expression levels increased in the biofilm mode of growth relative to planktonic cells over 2.0-fold for genes *AtzB* and *AtzC* but not at statistically significant levels. The remainder of the genes assayed, *AtzD, AtzE*, and *AtzF*, demonstrated moderate expression between 1.0 and 2.0-fold in biofilm mode of growth compared to the control planktonic condition of *Pseudomonas* sp. ADPT cells (FIG 4). The control condition is designated by a normalized expression of (+/-) 1.0 relative to increases (above abscissa) or decreases (below abscissa). Results and error under abscissa were calculated by taking the (-) inverse of the n-fold change to equate the represented bars. The expression of atrazine-degrading genes in biofilms relative to planktonic *Pseudomonas* sp. strain ADP cells were not found to be significantly different at an α = 0.05 (p = 0.2197).

**FIG 4.**
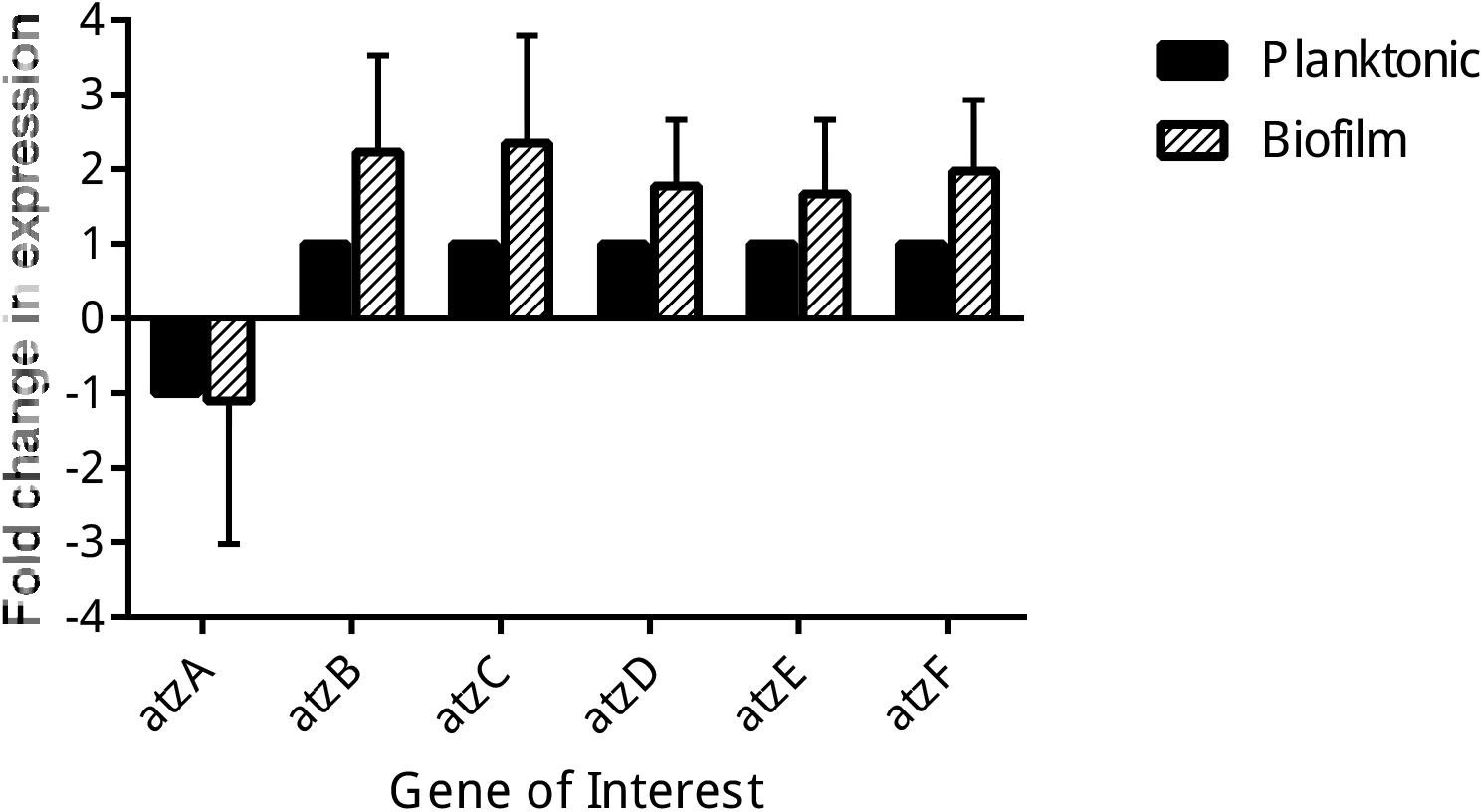
The fold change in gene expression of planktonic *Pseudomonas* sp. ADP cells (control) relative to *Pseudomonas* sp. ADP biofilms for each gene involved in the catabolic degradation of atrazine. The ordinate is scaled equally for downregulated (negative) and upregulated (positive) genes.

The atrazine-degrading genes assayed for *Pseudomonas* sp. strain ADP biofilms under varied temperature conditions demonstrated differential expression (FIG 5). For the first two atrazine-degrading genes, *AtzA* and *AtzB*, expression was decreased considerably for biofilms grown at 37°C relative to the control condition (30°C). *Pseudomonas* sp. strain ADP biofilms grown at 25°C, however, demonstrated relatively negligible changes in expression. Gene *AtzC* decreased in expression for both experimental temperature conditions, but more noticeably for 37°C grown biofilms. The three remaining genes, *AtzD, AtzE*, and *AtzF*, exhibited approximately 3.0-fold, 1.5-fold, and 3.0-fold increases in expression for biofilms grown at 37°C relative to the control condition, respectively. Minor increases in expression were also observed for 25°C for genes *AtzD* and *AtzE*, however, *AtzF* revealed over a 2.0-fold increase for the same growth temperature. The difference in expression of atrazine-degrading genes in *Pseudomonas* sp. strain ADP biofilms was not statistically significant between biofilms grown at 25°C and 30°C at an α = 0.05 (p = 0.8717), nor between biofilms grown at 37°C and 30°C (p =0.3748).

**FIG 5.**
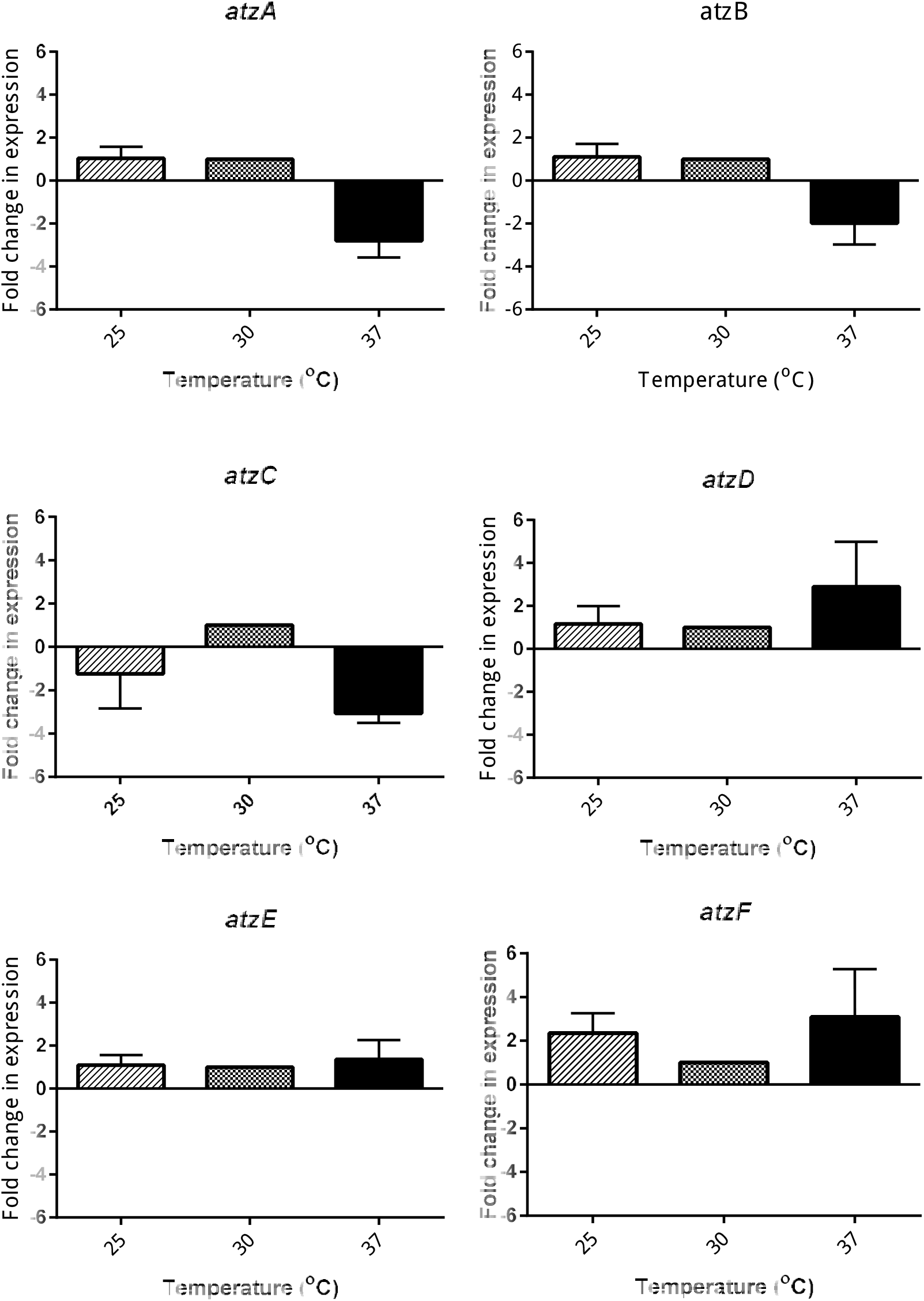
The fold change in gene expression of *Pseudomonas* sp. ADP biofilms grown at temperatures: 25°C, 30°C (control), and 37°C to maturity. Each gene involved in atrazine-degradation was assayed, and the ordinate scale is scaled for equal representation of upregulated (positive) and downregulated (negative) genes.

Gas chromatography-Mass spectrometry (GC/MS) was used to quantify atrazine concentration over five day periods in biofilm and planktonic cell modes of growth (FIG 6). Feed of the influent medium was 50 mg L^-1^. The atrazine effluent decreased linearly in biofilm-mediated cells until reaching an atrazine effluent of 20 mg L^-1^ at two days. The effluent subsequently increased from day two to day five, reaching a local maximum of 30 mg L^-1^. In shake flasks, the concentration decreased to 18 mg L^-1^ after 24 hours, followed by a constant level of atrazine until day four. The last 24 hours of planktonic cells demonstrated a slight decrease to 16 mg L^-1^.

**FIG 6.**
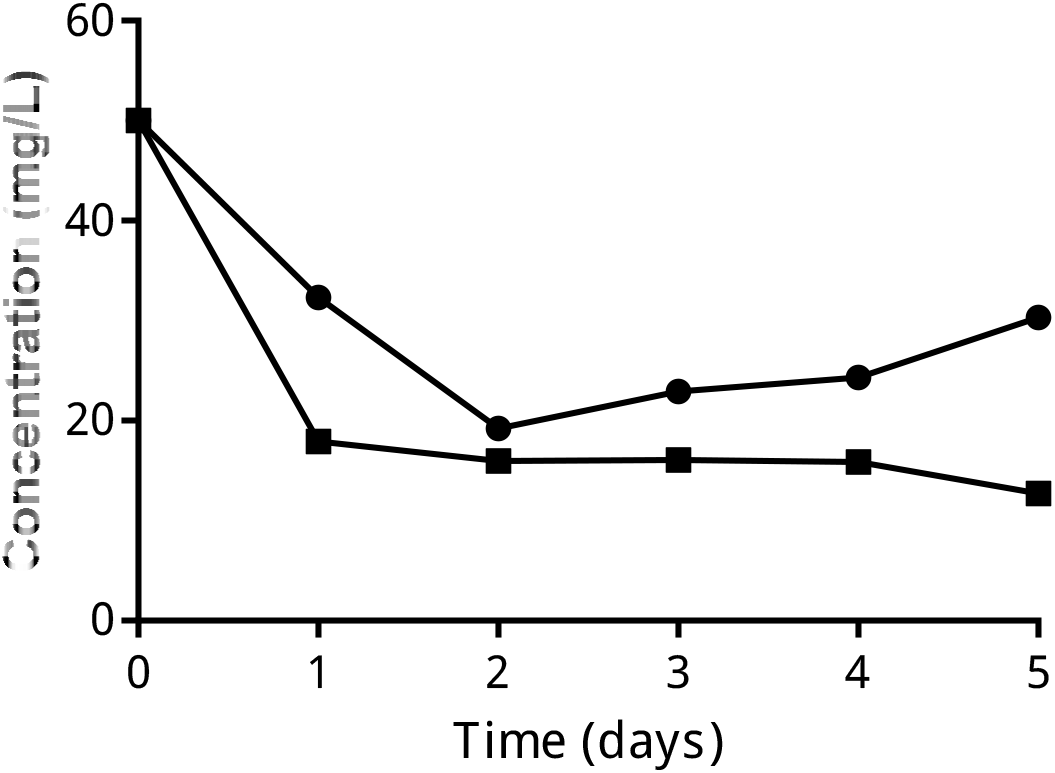
Atrazine concentration measured by GC-MS over five days in shake flasks containing a planktonic cell medium (◼) and a DFR-grown biofilm containing reactor effluent (•) as a result of microbial degradation.

The concentration of atrazine was measured in drip flow reactor effluent of *Pseudomonas* sp. strain ADP biofilms over a period of five days for two temperature conditions (30°C and 37°C) using GC/MS (FIG 7). Feed atrazine concentration was 30 mg L^-1^. The atrazine concentration for biofilm effluent at 30°C decreased linearly until 60 hours before reaching a minimum of 7 mg L^-1^, and then rising to 11 mg L^-1^ at five days. The level of atrazine in the biofilm effluent at a higher temperature condition, 37°C, initially increased from 21 mg L^-1^ to 26 mg L^-1^ at 38 hours, and then decreased considerably to 12 mg L^-1^ after 60 hours. From three days to five days, the level of atrazine decreased gradually to 6 mg L^-1^. Neither temperature condition resulted in undetectable atrazine levels in the effluent after five days of biofilm growth.

**FIG 7.**
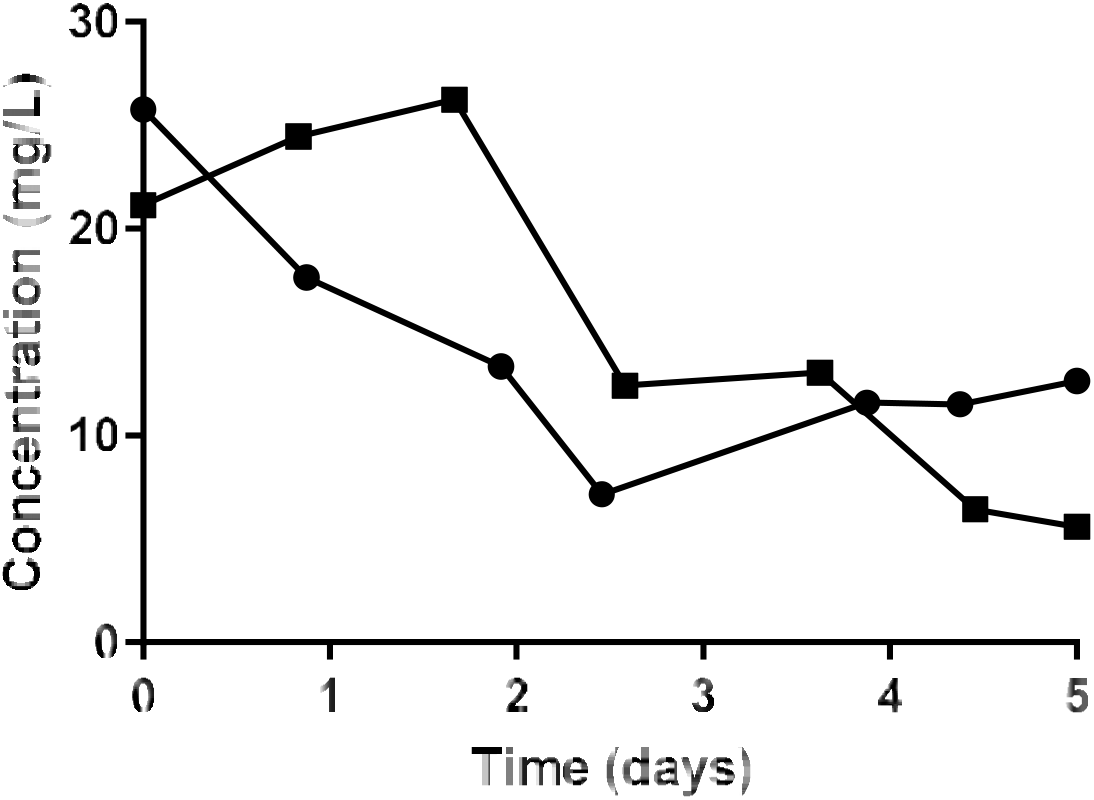
Atrazine concentration measured by GC-MS over five days in a DFR-grown biofilm (reactor effluent) at incubation temperatures of 30°C (•) and 37°C (◼) as a result of microbial degradation.

Micrographs elucidating surface morphology of planktonic cells and biofilms of *Pseudomonas* sp. strain ADP were captured with scanning electron microscopy (FIG 8). Both micrographs are cells grown in mineral medium containing atrazine (30 mg L^-1^) as the sole nitrogen source. Planktonic cells demonstrate a lack of growth networks and notable absence of any extracellular polymeric matrix, and cells in planktonic micrograph are observed to be rod-shaped, which is consistent and characteristic of Pseudomonad bacteria. The biofilm image captured with scanning electron microscopy exhibits a clear, complex network of strands and webbing intercalated between bacterial cells, forming an extracellular polymeric matrix which is remarkably characteristic of biofilm morphology. The scanning electron microscopy images can be visually differentiated based on mode of growth.

**FIG 8.**
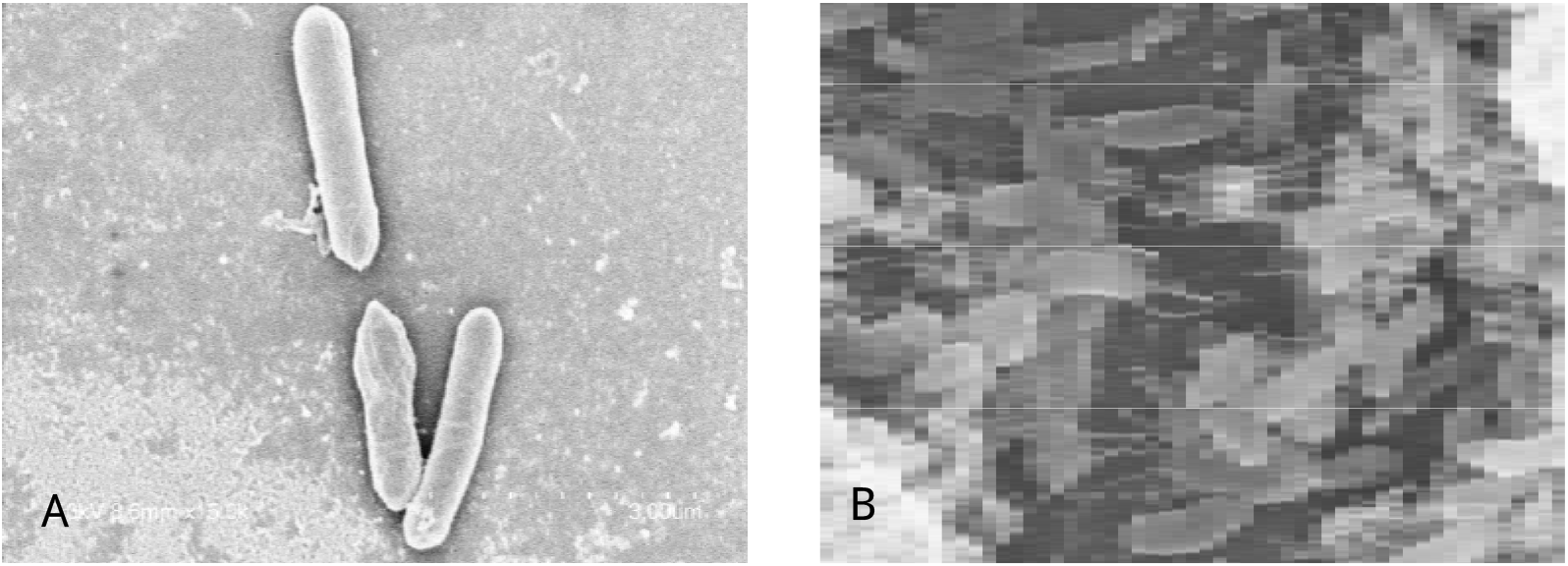
Scanning electron micrographs of planktonic *Pseudomonas* sp. ADP cells (A) and *Pseudomonas* sp. ADP biofilms (B) grown on minimal medium containing atrazine (30 mg L^-1^).

## Discussion

In our studies, *Pseudomonas* sp. strain ADP cells followed a prototypical growth curve when grown on atrazine or cyanuric acid as their sole nitrogen source. It has been previously reported that *Pseudomonas* sp. strain ADP could utilize a metabolite of its catabolic pathway, cyanuric acid, as source for growth and survival.(26–28) The bacterial isolate has also demonstrated growth on the synthetic herbicide, atrazine.(10) The increase is the rate of cell proliferation for atrazine-grown bacteria relative to cyanuric acid-grown bacteria may demonstrate the plasmid, pADP-1, containing the genes encoding for the catabolic enzymes, is more efficient for cellular growth when the cells are able to utilize the initial substrates on the regulated pathway.

Following from the results of the gene expression experiments, we chose to employ relative expression instead of quantitative expression as we are interested in comparing two modes of growth. This experiment has been performed to examine differences in gene expression between *Salmonella typhimurium* and *Staphylococcus aureus*, which aided in elucidating expression patterns of antibiotic-resistant pathogens.(29) A method previously developed and optimized by Devers regarding reverse transcription quantitative real-time PCR (RT-qPCR) for *Pseudomonas* sp. strain ADP cells was used for these studies.(13) The negligible decrease in the expression of *atzA* in biofilms relative to free cells suggests atrazine disappearance is caused by mechanisms other than metabolic processes. The extracellular polymeric substances of the biofilm may cause atrazine to be entrapped within the matrix by a mechanism of biosorption, thereby removal of atrazine by a non-catabolic process. There is no sufficient evidence to suggest that there are significant changes in expression of atrazine-degrading genes and consequently increased or decreased catabolic activity of biofilm-associated cells with regards to contaminant degradation based on mRNA levels alone. These results demonstrate despite similar rates of metabolism in planktonic cells and biofilms, atrazine also undergoes remediation by a physical process such as adsorption based on gas chromatographic results.

In concurrence, the atrazine disappearance was monitored in both the suspended cell and biofilm modes of growth. The relatively constant decrease in atrazine in free-cells over time in consistent with previous work by Devers, and indicates the contaminant may follow first-order kinetic degradation patterns.(13) However, compared to batch systems (suspended cells grown in shake flasks), flow-systems (biofilms grown in drip-flow reactors) reveal increasingly complex degradation kinetics. Initially the rapid decrease in atrazine would likely be due to the contaminant entrapment within the extracellular polymeric matrix while the substrate is being utilized by biofilm-enclosed cells, as evidenced in previous studies by Henry, Jessop, and Peeples.(18) The increase in atrazine concentration in the biofilm flow system likely results from dispersal of biofilm-mediated cell release into the effluent after maturation. Following maturation, we would expect the biofilm to degrade atrazine at a greater efficiency in a period from five to ten days of growth. Due to the complex nature of biofilms, including proliferation of a self-produced extracellular matrix, close cellular proximity, increased levels of quorum sensing, differential nutrient bioavailability and exchange of genetic material all contribute to the remediation potential of biofilms for degrading recalcitrant pollutants.(20)

In the natural environment, temperature is varied according to season, and for industrial systems, the temperature of biofilm reactors can be optimized for degradation processes. Previously, it has been reported that for every rise in 10°C, the rate of degradation of the pollutant toluene increased nearly 2-fold in soil and water.(30, 31) In *Pseudomonas* sp. strain ADP biofilms, decreasing the temperature from 30°C to 25°C resulted in no differential expression and from a genetic perspective, has no increase or decrease in functional gene products that may exhibit higher or lower catabolic potential. After increasing the temperature of biofilm growth to 37°C, the decreased expression of the initial three genes (*AtzA, AtzB*, and *AtzC*) could suggest either I) a decrease in catabolic activity of transformations of atrazine to cyanuric acid II) loss of genes *AtzA, AtzB*, and *AtzC* as they have greater instability on the plasmid pADP-1 compared to the latter genes in the pathway, or III) increased temperature causes pathway regulatory elements for the initial three genes to decrease in overall mRNA levels. Interestingly, the increase in expression of the remaining three genes (*AtzD, AtzE*, and *AtzF*) could help elucidate the regulation of the catabolic pathway with regards the cyanuric acid degradation. If the temperature of biofilm growth is increased to 37°C, this could ensure stability of the operon-like gene set *AtzDEF* on the plasmid, pADP-1 and degrading the intermediates from cyanuric acid to allophanate at increased rates from higher enzymatic activity.

After monitoring atrazine degradation, for 30°C, the control, and 37°C, in *Pseudomonas* sp. strain ADP biofilms, after a period of five days both resulted in the disappearance of atrazine to 12 ppm (30°C) and 5 ppm (37°C). The characteristic increase in the higher temperature condition, similar to the previous biofilm degradation curve, may be a result of atrazine trapped within the extracellular polymeric substances, followed by degradation and release. At the end of monitoring period and growth of the biofilms, a greater proportion of atrazine was not detectable by GC/MS for the biofilm grown at 37°C. This phenomenon could be the basis for design of reactor systems with temperature-controlled remediation. Furthermore, in the state of Iowa and Illinois where atrazine contamination is prevalent, biofilms forming the surrounding community that include *Pseudomonas* sp. strain ADP may be able to more effectively degrade atrazine in seasons where the local temperature of the microenvironment increases to 37°C.(31) This may occur, for example, in Iowa where near-surface soil conditions can reach average temperatures from 25°C to 33°C in mid-summer.(32)

Scanning electron micrographs of biofilms appeared to have an intricate structure of webbed material surrounding cells, further confirming biofilm growth at five-days. Similar studies by Henry, Jessop, and Peeples have validated the appearance of webbed structures as a component of the extracellular polymeric substances in *Pseudomonas* sp. strain ADP biofilms.(18) The free cells, in contrast, are lacking any networking between individual cells grown in the mineral medium.

Numerous experiments have been conducted elucidating gene expression, isolation of enzymes, and degradation kinetics of atrazine in *Pseudomonas* sp. strain ADP suspended cells, while relatively few experiments have been conducted on expression techniques and degradation kinetics on biofilms of the same type.(33–37) Despite similar catabolic activity between atrazine-degrading genes in planktonic and biofilm cells, the key result demonstrates the immense potential of employing biofilm-based reactors for *ex situ* bioremediation through processes such as biosorption via the EPS and exploring the complex mechanisms behind natural films formed in the environment for *in situ* remediation technologies. Moving forward we will develop a method to visualize the expression of atrazine-degrading genes in *Pseudomonas* sp. strain ADP biofilms using confocal microscopy, and further generalize the method to apply across strains with contaminant remediation potential. By doing so, we will be able to also monitor the transient appearance and disappearance of intermediates involved in several pathways of degradation. Another future area of interest will be to further analyze gene expression in *Pseudomonas* sp. strain ADP biofilms grown on environmentally and industrially-relevant intermediates, such as cyanuric acid, to evaluate the films’ biodegradation efficiency on related compounds.

## Conclusion

The objective of this work was to elucidate the expression of the atrazine-degrading gene set in *Pseudomonas* sp. strain ADP cells based on mode of growth in a sessile state as a biofilm relative to suspended cells grown in shake flasks. After evaluating the complete gene set of interest in *Pseudomonas* sp. strain ADP, no significant differences were quantified in the expression of atrazine-degrading genes in biofilm-mediated cells relative to suspended cells grown on atrazine. When biofilms were grown at higher temperatures, an increase in the expression of genes *AtzD, AtzE*, and *AtzF* may designate increased degradation from the metabolite cyanuric acid to carbon dioxide and ammonia. The degradation patterns of atrazine differed in the biofilm and suspended cell state based on GC/MS analysis due the unique structures and cellular activity of biofilm growth, which is substantiated by scanning electron micrographs. The gene expression and kinetic biofilm data provided here will aid in the development of biofilm technologies to assist in bioremediation of recalcitrant herbicides, such as atrazine. Our future studies would aim to include analyzing the kinetics in real-time of each pathway metabolite in the biofilms and suspended cells. Additionally, evaluating the expression of genes in biofilms via the same methodology presented here may be used for genetic analysis in medically-relevant biofilms such as the formation of films on medical devices and their presence in lung disease. Furthermore, the results here can be generalized to demonstrate how gene expression results and analytical chemical data can synergistically provide greater insight into bioremediation and microbial degradation.

## Materials and Methods

### Bacterial strain, cultivation medium, and culture conditions

A frozen glycerol stock of *Pseudomonas* sp. strain ADP (DSM 11735) was streaked onto an agar plate containing mineral medium with 1000 mg L^-1^ atrazine (3.50 g L^-1^ Na_2_HPO_4_ • 2H_2_O, 1.00 g L^-1^ KH2PO4, 0.10 g L^-1^ MgCl2 • 6H2O, 0.05 g L^-1^ CaCl2, 1 g L^-1^ d-glucose, 1.00 mL L^-1^ (v/v) trace element solution, 15 g L^-1^ noble agar, pH 7.25). The streak plate was incubated overnight (30°C) and a single colony was inoculated in a liquid mineral medium containing 30 mg L^-1^ atrazine (3.50 g L^-1^ Na2HPO4 × 2H2O, 1.00 g L^-1^ KH2PO4, 0.10 g L^-1^ MgCl2 × 6H2O, 0.05 g L^-1^ CaCl_2_, 1 g L^-1^ d-glucose, 1.00 mL L^-1^ (v/v) trace element solution, pH 7.25) in shake flasks (125 mL) at 30°C. The agitation rate was maintained at 250 rpm. After 12 hours, cell growth was measured indirectly using spectrophotometry detailed in the proceeding section. If growth was sufficient in the pre-inoculum (OD_600_ > 0.1), the culture was passed to fresh medium (50 mL) in a larger flask (250 mL) while maintaining a 10% inoculum.

### Growth of Pseudomonas sp. strain ADP suspended cells

Cellular growth of the inoculum was measured indirectly using spectrophotometry (Thermo Spectronic BioMate, Waltham, MA). Optical density measurements (OD_600_) were taken once per hour for 100 hours from second pass of cells. Samples were then diluted ten-fold when OD_600_ > 0. 3 to remain within the instrument’s limit of detection. The growth curve was used to determine the relative growth rate, doubling time, and time to reach exponential phase. Overnight cells were passed to a larger inoculum and samples were collected at previously determined exponential phase (t = 30 hours) for consistency during RNA extraction and microscopy experiments.

### Drip-flow reactor for Pseudomonas sp. strain ADP biofilm growth

Biofilms were grown by the method of Tolker-Nielsen in a drip-flow reactor with minor modifications.(38) A four-channel Teflon Drip Flow Reactor (10 cm × 2.5 cm × 2 cm) was used to grow *Pseudomonas* sp. strain ADP biofilms. Briefly, cell cultures in exponential phase (OD_600_ > 0.2) from shake flasks were harvested by centrifugation (6000 x g, 10 minutes, 4°C) and resuspended in mineral medium (10 mL) containing atrazine (30 mg L^-1^). Re-suspended cells were dispersed on Silane-coated microscope slides and incubated at 30°C for 6-8 hours in batch mode to allow for bacterial attachment. Mineral medium containing atrazine (30 mg L^-1^) was transmitted by peristaltic pumps (rate of 0.8 mL minute^-1^) to flow over the slide surface with adhered bacteria until the biofilm reaches maturation (five days) (FIG 2).

### Temperature dependence and mode of growth studies

Biofilms were grown in the drip-flow reactor during continuous phase at three temperatures (25°C, 30°C, 37°C) in a controlled incubator to determine the effect of temperature on mRNA levels of atrazine-degrading genes using RT-qPCR. Congruently, bacterial biofilms were harvested at full maturation (5 days) and planktonic cultures were harvested at exponential phase (30 hours) and subjected to an RT-qPCR workflow to determine the effect of mode of growth on mRNA levels of atrazine-degrading genes.

### RNA extraction and quantification

Optimization of extraction for RNA biofilms was achieved by modifications of two established methods.(39, 40) Both suspended cells and biofilms were subjected to the same extraction method to achieve consistency. Briefly, biofilms were scraped from each slide into centrifuge tubes (50 mL) containing Trizol reagent (2 mL). Biofilm suspensions and planktonic samples were vortexed briefly, then sonicated using a sterile probe for 10 seconds at medium-high output (pulse: 4/10 power, Sonic Dismembrator 550, Fisher Scientific, Waltham, MA). The insoluble material of each sample was removed by centrifugation (6,000 × g, 10 minutes, 4°C). The supernatant containing the RNA was extracted with the remaining Trizol according to the manufacturer’s instructions. RNA precipitated from the isolation procedure was purified and DNase-treated using spin column purification (Qiagen RNeasy Mini Kit, Germany) and on-column enzyme treatment (Qiagen DNase-set, Germany), respectively. Bacterial RNA was quantified and measured for integrity using micro-spectrophotometry (Thermo NanoDrop, Waltham, MA) and capillary electrophoresis (Agilent Bioanalyzer, Santa Clara, CA). Samples containing RNA greater than 50 ng μL^-1^ and an RNA integrity number above 8.0 were used for reverse transcription reactions.

### Reverse transcription

RNA samples were diluted to approximately 50 ng μL^-1^ before proceeding with reverse transcription. Samples were reverse transcribed using a high-capacity reverse transcription kit (Thermo Fisher, Waltham, MA). Briefly, 2 × RT master mix (10 μL) and the RNA sample (10 μL) were added to PCR tubes, mixed, and placed in a thermal cycler under the following conditions: 10 minutes at 25°C, 120 minutes at 37°C, and 5 minutes at 85°C. Complementary DNA was stored at −20°C until quantitative PCR step. Samples containing kit components in the absence of the enzyme were used as negative controls.

### Quantitative PCR

Relative quantification RT-qPCR was chosen for the methodology to determine fold changes in expression within groups of temperature dependence and mode of growth. Validation experiments were performed for each primer pair of the target and reference gene to ensure efficiencies are approximately equal. The qPCR assays were the method of Devers with minor modifications.(13) For qPCR, 25 μL reactions were prepared in 96-well plates containing: 12.5 μL Power SYBR Green Master Mix (Thermo Fisher, Waltham, MA), 2.0 μL forward primer (10 μM), 2.0 μL reverse primer (10 μM), 5-fold diluted cDNA sample (12.5 ng), and nuclease-free water. No template controls were added to each plate. Each biological replicate (3×), reverse transcription replicates (2×), and technical replicates (3×) were run under the following qPCR conditions: 10 minutes at 95°C for enzyme activation, and 40 cycles of 15 seconds at 95°C, 15°C at optimal hybridization temperature, and 15 seconds at 72°C.(13) Dissociation curves were performed for all qPCR assays. All RT-qPCR analysis was performed using ABI software. (Applied Biosystems ABI SDS version 2.4.1) The ΔΔCt method was used to calculate the fold-difference between the calibration samples and experimental samples. In the temperature dependence group, the central temperature condition 30°C was chosen as the calibration sample. In the mode of growth group, the suspended cells were chosen as the calibration sample. The ΔCt values were calculated between the reference gene (16s *rDNA*) and the gene of interest (*AtzA, AtzB, AtzC, AtzD, AtzE, AtzF*). All calibration samples were normalized to −1 and +1 to calculate the relative fold change in test conditions.

### Analytical methods

GC/MS (Waters GCT Premier, Milford, MA) was used to determine the concentration of atrazine in biofilm and suspended cell samples. In the biofilm condition, samples (1.5 mL) were collected in triplicate from the bioreactor effluent every 18-24 hours for five days. In the suspended cell condition, samples (1.5 mL) were retrieved in triplicate from incubated shake flasks (30°C, 250 rpm) every 18-24 hours for five days. For temperature-controlled biofilms, samples (1.5 mL) were collected in triplicate (30°C, 37°C) from the bioreactor effluent every 18-24 hours for 5 days. Atrazine was extracted from the effluent with ethyl acetate following slight modifications of an established protocol.(41) Organic material and atrazine were separated from the aqueous phase by adding ethyl acetate (0.5 mL) to effluent (1.0 mL) and spinning briefly. The upper organic phase was drawn off and transferred to a micro centrifuge tube. The bottom aqueous phase was extracted twice using ethyl acetate and the organic fractions were combined in a micro centrifuge tube. The extracted organic phase samples containing atrazine were dried using centrifugal evaporation (Thermo Fischer Savant SpeedVac, Waltham, MA). Dried extracts were re-suspended in ethyl acetate (1 mL), vortexed, and transferred to clear vials. Terbuthylazine, a compound within the s-triazine class of herbicides, was added as an internal standard due to its structural similarity to atrazine and absence in the bacterial degradation pathway.(42) A calibration curve was established using concentrations of atrazine from 1 mg L^-1^ to 60 mg L^-1^.

### Scanning Electron Microscopy

Biofilms harvested after five days and planktonic cells harvest at mid-logarithmic phase on mineral medium containing atrazine were prepared for scanning electron microscopy. Biofilms were grown on cover slips, while cells in suspension were transferred to micro tubes. In both conditions, cells were fixed in 2.5% glutaraldehyde, rinsed with cacodylate buffer thrice in an hour (3 × 20 minutes). Subsequently cells were subjected to 1% osmium tetroxide in buffer for two hours, followed by another rinse in cacodylate buffer thrice in an hour (3 × 20 minutes). Cells were rinsed with deionized water and subjected to an ethanol dehydration series (15-30 minutes in each solution of 25%, 50%, 75%, 95%, and 100% ethanol). Hexamethyldisilane was used as a substitute for critical point drying. Biofilms and suspended cells were then mounted on stubs, sputter coated, and examined in the scanning electron microscope.

### Statistical Analysis

All gene expression studies were subjected to a two-tailed student’s t-test to examine n-fold changes in expression for significance at an alpha of 0.05 with n of 3. Statistical analyses were carried out following propagation of error and final fold change values. Tests were performed to determine significance of fold change in expression of atrazine-degrading genes between biofilm grown cells and planktonic grown cells. Additionally, tests were performed to determine significance of fold change in expression of atrazine-degrading genes between biofilms grown at higher (37°) and lower (25°C) temperatures relative to a control (30°C). Lastly, tests were performed to determine significance of fold change in expression of three atrazine-degrading genes (*AtzD, AtzE*, & *AtzF*) in cells grown on mineral medium containing cyanuric acid relative to a control of cells grown on mineral medium containing atrazine.

## Acknowledgments

The authors would like to acknowledge the National Institutes of Health and Center for Biocatalysis and Bioprocessing for the training grant that made this work possible, the Genomics Division at the University of Iowa for experimental design considerations, Dr. Lawrence Wackett’s Research Group for providing streak plates containing the strain *Pseudomonas* sp. strain ADP, and Dr. Marion Devers for all correspondence relating to research on strain *Pseudomonas* sp. strain ADP. The funders had no role in study design, data collection and interpretation, or decision to submit the work for publication. The authors declare no conflict of interest, financial or otherwise.

